# An early surge of norepinephrine along brainstem pathways drives sensory-evoked awakening

**DOI:** 10.1101/2025.01.29.635485

**Authors:** Noa Matosevich, Noa Regev, Uddi Kimchy, Noam Zelinger, Sina Kabaha, Noam Gabay, Amit Marmelshtein, Yuval Nir

## Abstract

The locus coeruleus norepinephrine (LC-NE) system regulates arousal and awakening; however, it remains unclear whether the LC does this in a global or circuit specific manner. We hypothesized that sensory-evoked awakenings are predominantly regulated by specific LC-NE efferent pathways. Anatomical, physiological, and functional modularities of LC-NE pathways involving the mouse basal forebrain (BF) and pontine reticular nucleus (PRN) were tested. We found partial anatomical segregation between the LC->PRN and LC->BF circuits. Extracellular NE dynamics in BF and PRN exhibited distinct sound-evoked activation during sleep, including a fast sound-evoked NE peak specific to PRN. Causal optogenetic interrogation of LC efferent pathways, by retro-ChR2 activation or PdCO silencing of synapses in target regions, revealed a pivotal role for early LC->PRN activity in driving arousal and sound-evoked awakenings. Together, our results uncover a prominent role for early LC-NE PRN activity in connecting sensory and arousal pathways and establish LC heterogeneity in regulating arousal.

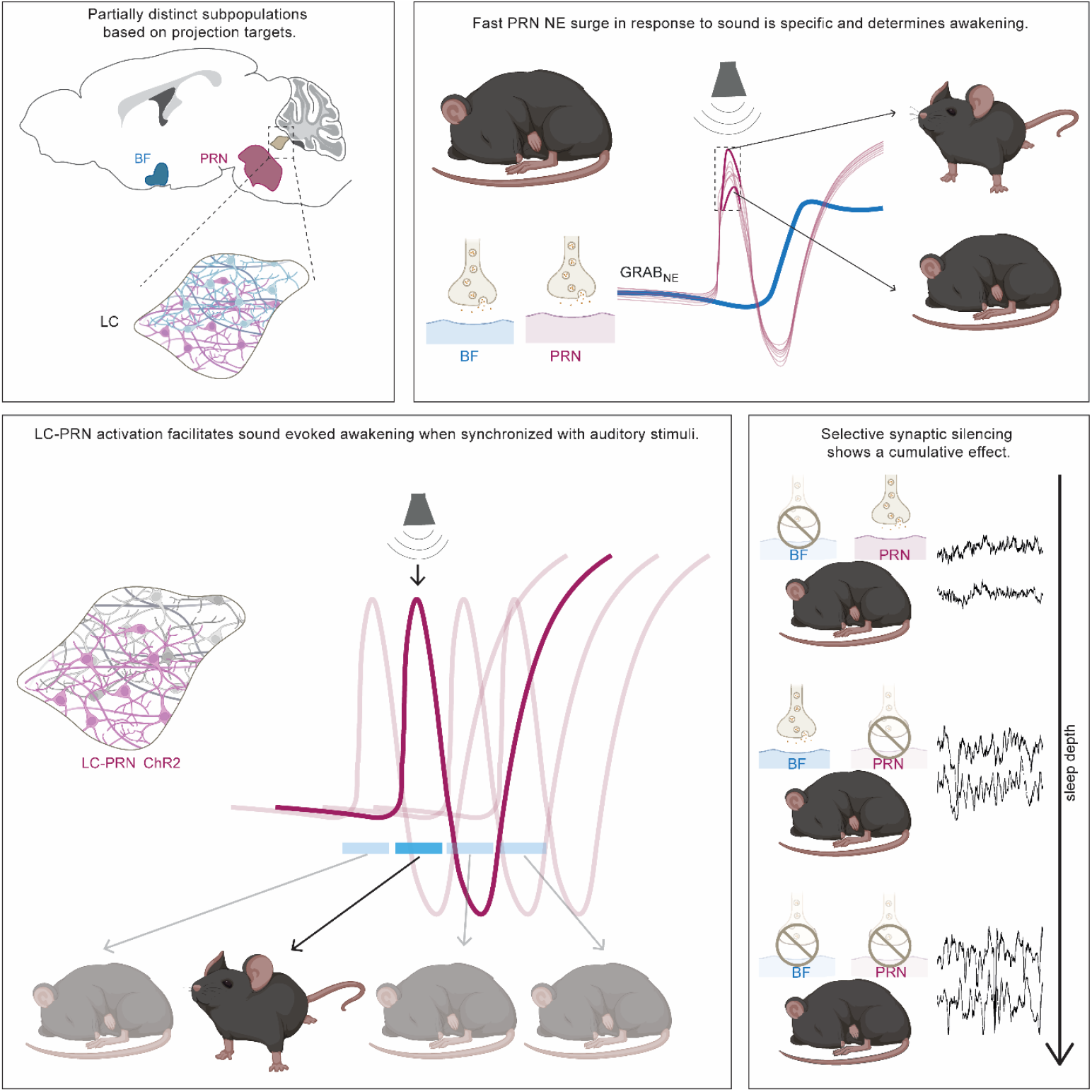

## Introduction

Engagement with the environment is critical to our survival as it allows us to perceive changes, respond to cues, and adapt behavior to diverse situations including potential threats. Sleep is dominated by reduced responsiveness to the environment, rendering animals vulnerable to predation. An elevated ‘arousal threshold’ characterizes sleep across all species and constitutes the main criterion by which sleep is defined in invertebrates lacking cortical EEG^1–5^, and yet the pathways underlying the neural basis of sensory disconnection during sleep are poorly understood.

An important neuromodulatory element in shaping responsiveness to external sensory events across internal states is locus coeruleus – norepinephrine (LC-NE) activity. Accordingly, LC-NE activity correlates with sleep fragility and micro arousals, and is inversely related to spindle activity associated with sleep continuity^6–8^. In wakefulness, LC-NE activity supports orienting responses towards behaviorally meaningful salient stimuli^9–13^. Moreover, LC-NE activity was recently shown to be a major determinant of sound-evoked awakening (SEA) in rats^14^. LC activation around auditory stimulation in sleep increased the probability of waking up in response to a sound, whereas LC silencing showed the inverse effect.

However, while LC-NE neuromodulation was traditionally regarded as globally homogenous^15–20^, recent studies identified some segregation of efferent LC pathways along with associated functional diversity (reviewed in^21^). For example, spinally-projecting LC neurons exert analgesic actions, whereas ascending LC projections show a pro-nociceptive effect^22–24^. LC neurons projecting to the prefrontal vs. motor cortex differ in their molecular profiles, excitability, and activity across vigilance states^25^. Furthermore, LC neuronal firing has been shown to be only sparsely synchronized^26^. This raises the question: could heterogenous LC-NE activity also play a role within the domain of arousal and sleep awakenings?

To address this, we investigated whether the effect of LC on SEA is mediated by distinct LC projections. We focused on two target regions, the pontine reticular nucleus (PRN) and the basal forebrain (BF) for representing brainstem and forebrain target regions, respectively, as both areas are profoundly innervated by LC-NE neurons and are implicated in regulating arousal^27–33^. We combined anatomical mapping of PRN or BF projecting neurons, GRAB_NE_-based extracellular NE recordings at these target regions, and bi-directional optogenetic manipulations of presynaptic LC terminals in these projections^34^. Our results reveal a specific role for an early surge of brainstem NE in driving sensory-evoked awakening from sleep.

## Results

### LC neurons projecting to the BF and PRN are partially distinct anatomically

We first investigated whether LC cells projecting to the BF and those projecting to the PRN exhibit anatomical segregation. Using retrograde tracing (Fig. 1A), retro-beads were injected either in the BF or the PRN (n=7 and n=8 respectively) of WT mice and successful targeting was histologically confirmed (Fig. 1B). LC subpopulation analysis (Fig. 1C) determined the distribution of LC neurons along the dorso-ventral (DV) and the anterior-posterior (AP) axes of the LC (Fig. 1D). We found a statistically significant difference in the anatomical distribution of BF-projecting vs. PRN-projecting LC neurons (p=0.004, Monte Carlo permutation test), implying that the BF and PRN neuronal projections of the LC are different subpopulations. Specifically, BF-projecting LC neurons were localized more dorsally in the LC compared to the PRN-projecting LC neurons (p=6e-04, Monte Carlo permutation test). No projection bias was observed along the AP axis (p=0.4, Monte Carlo permutation test).

**Figure 1:**
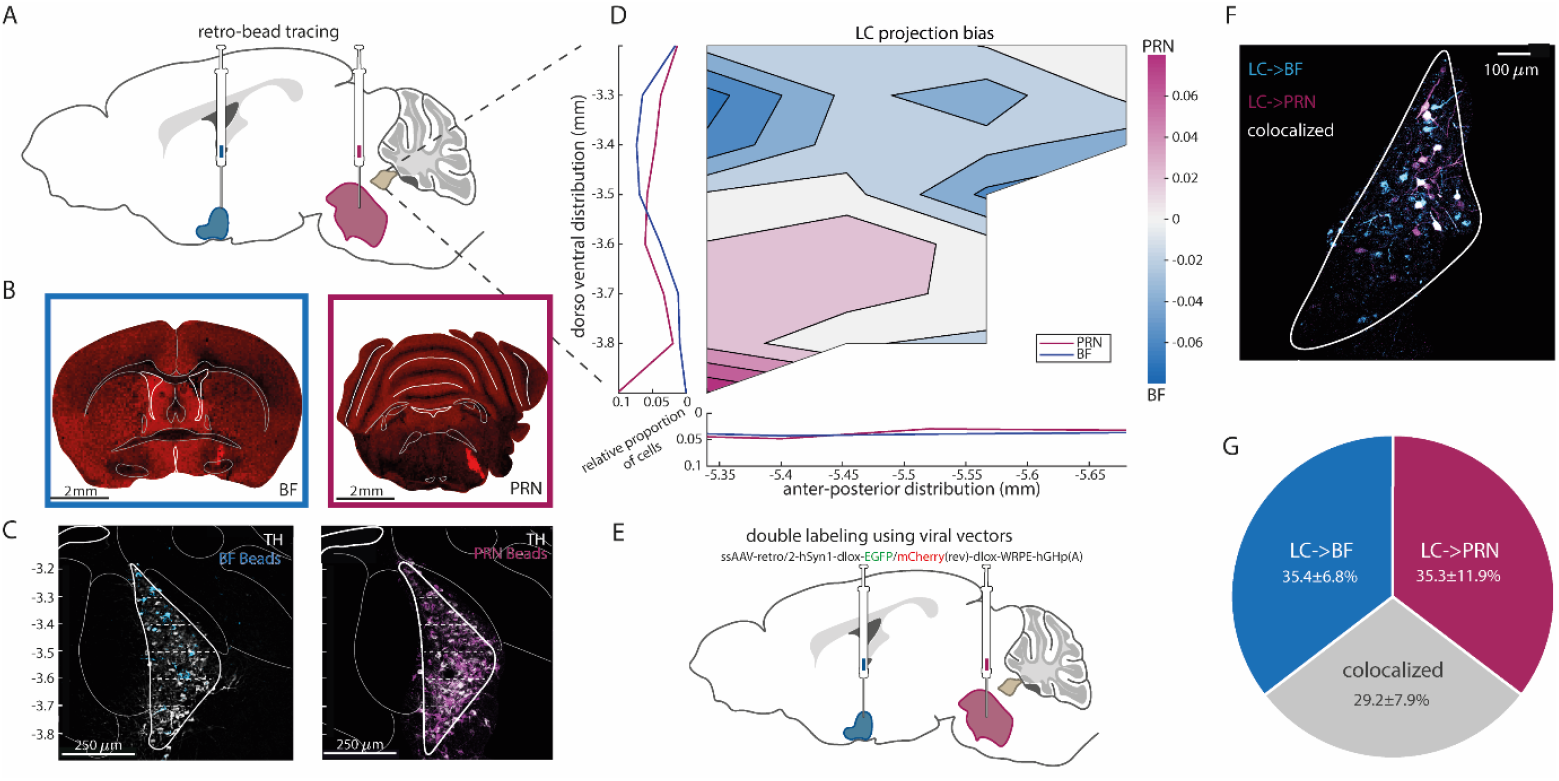
LC neurons projecting to the BF and PRN are partially distinct anatomically. **A**, Surgical approach-sagittal section taken from Allen Mouse Brain Atlas. **B**, Representative examples of BF retro-bead injection (left) and PRN retro-bead injection (right). **C**, The LC section at -5.4 mm from bregma (AP). BF bead tracing in blue (left), PRN bead tracing in magenta (right), tyrosine hydroxylase (TH) staining in white. A mask was applied to highlight the LC region specifically since retrobeads are not specific in their tagging. **D**, Normalized projection pattern of LC neurons to BF or PRN calculated as 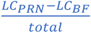 calculated over n=8 PRN injected mice and n=7 BF injected mice (see methods). Blue tones on the heatmap represent a higher probability of BF projecting cells, and magenta tones represent a higher probability of PRN projecting cells. The image portrays the LC in a sagittal section. To the left are the average traces of the LC cell distribution on the dorso-ventral (DV) axis from the top corner of the LC (-3.2 mm relative to bregma) to -3.9 mm from bregma. At the bottom are the average distributions of the LC neurons on the antero-posterior (AP) axis from -5.34 mm to -5.7 mm from Bregma. Sections were identified based on the Mouse Brain Atlas^35^ and thus coordinates were determined. **E**, Sagittal section taken from Allen Mouse Brain Atlas to portray the approach. **F**, LC section of representative mouse. BF-blue, PRN - magenta, white-colocalized cells. **G**, Pie chart of colocalization percentage. Average and standard deviation calculated over n=5 DBH-Cre mice.

Next, to determine the degree of cellular co-localization among the two subpopulations, we conducted retrograde viral tracing by performing simultaneous injections of red and green Cre-dependent AAVretro viruses into both PRN and BF (n=5 DBH-Cre mice, Fig. 1E,F). The quantitative analysis revealed that tagged LC neurons consisted of 35.34 ± 11.95% PRN-only projecting cells, 35.44±6.79% BF-only projecting cells, and 29.21±7.93% co-localized cells (projecting to both target regions), meaning that the LC contains both distinct subpopulation as well as neurons sending axonal collaterals to both BF and PRN. T-tests against the null hypothesis of homogeneity was significant, confirming that each subpopulation is distinct compared to the overlapping subpopulation, and that neither LC-> PRN nor LC-> BF is a subpopulation of one another. (LC->PRN: p=0.0041, t(4)=5.91; LC->BF: p=4.669e^-04^, t(4)=10.49; Fig 1G). Thus, beyond a dorsal-ventral gradient in the distribution of BF-projecting LC cells, most tagged LC neurons project either to PRN or to BF but not to both targets, establishing anatomical modularity in brainstem vs. forebrain LC projections.

### A fast brainstem-specific auditory-evoked NE response predicts awakening

Next, we compared extracellular NE dynamics between brainstem and forebrain target regions with GRAB_NE_^36^. We first validated our GRAB_NE_ tool by testing BF GRAB_NE_ dynamics in response to optogenetic LC stimulation (Fig S1). Results revealed a dose-dependent increase in pupil dilation and NE levels with stronger optogenetic LC stimulation in lightly anesthetized mice^14^. In addition, induced sleep-to-wake transitions in freely behaving mice were associated with elevated NE levels and EEG activation^37^, confirming that GRAB_NE_ records NE dynamics that are controlled by LC neurons.

To directly compare between NE dynamics in the BF and PRN, WT mice (n=7) were injected with GRAB_NE_ in both BF and PRN and fiber photometry was recorded simultaneously from the two areas during natural sleep (Fig 2A,B). Around spontaneous awakenings from non-rapid eye movement (NREM) sleep (Fig S2), both target regions exhibited similar dynamics of increased NE levels with no discernable difference between the two (p=0.53, t(6)=0.66; Fig 2C). By contrast, sound-evoked awakening (SEA) experiments (Fig 2D) revealed distinct NE dynamics in the PRN compared with the BF (Fig 2E). While the PRN NE dynamics exhibited a fast surge in response to sounds during sleep, BF NE showed a slower, later response. Experiments with a mutated GRABconstruct confirmed that the early positive peak indeed represents NE dynamics rather than an artifact (Fig 2G). Time series analysis across mice (STAR Methods, Fig 2F) revealed a time window in the PRN NE trace where activity was significantly different than baseline: ‘PRN surge’ (t=0.3-0.6s). Similar analysis in the BF NE traces revealed a different time window with significant activity: ‘BF rise’ (t=1.8-3s).

**Figure 2:**
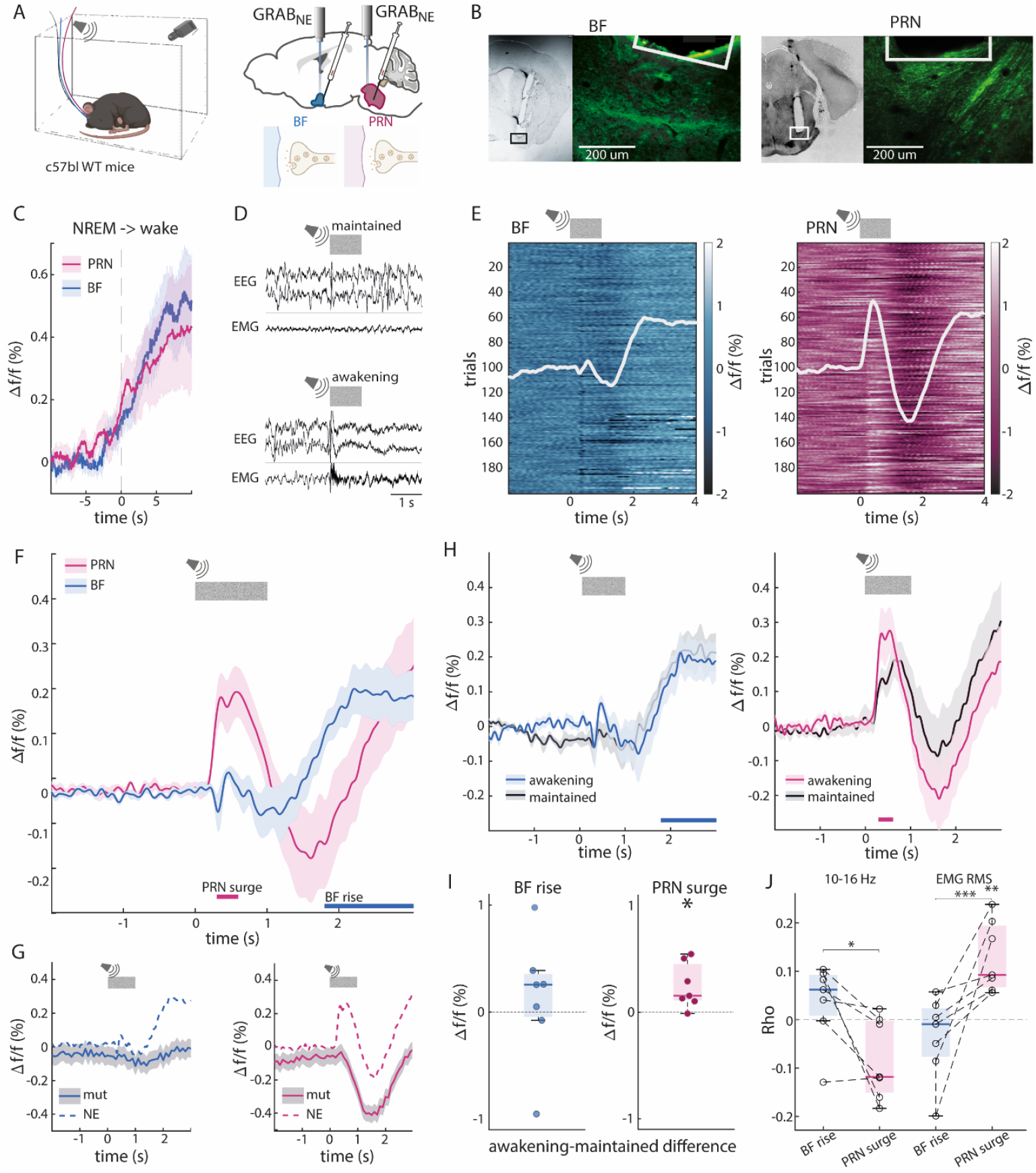
A fast brainstem-specific auditory-evoked NE response predicts awakening. **A**, Depiction of experimental setup (left), and of surgical approach (right). **B**, Representative examples of fiber and GRAB_NE_ expression in BF (left) and PRN (right). Green-GFP. **C**, Average GRAB_NE_ traces in BF (blue) and PRN (magenta) during spontaneous awakening from NREM sleep across mice (n=7). **D**, Top-depiction of sound evoked arousal (SEA) experiment: broadband noise was played in 3 intensities (see methods). Each sound was played for 1 sec with 59 s ± 0.5 s jitter. Bottom-examples of trials that led to maintained sleep (middle) or awakening (bottom). **E**, All NREM SEA trials in a representative mouse. Left – BF, right-PRN. White graph overlaid represents the average trace. **F**, Average NREM SEA traces in BF (blue) and PRN (magenta) across mice (n=7). Horizontal lines represent time periods that are significantly above 0 for PRN and BF in magenta and blue respectively. **G**, As a control group, GRAB_mut_ experiments were run on n=4 mice in BF (blue) and n=4 mice in PRN (magenta). GRAB_mut_ experiments were conducted on mice with only one injection site and fiber placement, either BF or PRN. The SEA experiment was run, and data was averaged across all trials forgoing scoring, but other than that using the exact same analysis pipeline as for GRAB_NE_. Dashed lines represent the corresponding traces of GRAB_NE_. **H**, Average traces across mice of maintained and awakening trials. BF-left, blue; PRN- right, magenta. Horizontal lines indicate BF rise and PRN surge respectively, which were the times significantly above baseline. **I**, Box plots of signal differences between trials with sound evoked awakening and trials with maintained sleep during BF rise and PRN surge. Left-BF rise, right-PRN surge. Dots represent single animals **J**, Correlation values between mean EEG and EMG and response elements (BF rise-blue, PRN surge-magenta). EEG 10-16 Hz band-left, EMG root mean square (RMS)-right. See methods for further detail on EEG power band analysis. EMG RMS was calculated over the trial (-5:15) EMG segment by binning to 100 ms and calculating for each bin the RMS. To account for variability across trials and animals, each trial was baseline subtracted. The 4 s from sound onset were averaged in each mouse.

Next, we examined which response element was associated with behavioral trial outcome (awakening vs. maintained sleep; Fig 2H,I). The early brainstem NE response (‘PRN surge’) was elevated in trials in which the sound elicited awakening compared to trials in which the mouse maintained sleep after sound (PRN surge: p=0.02, t(6)=-3.06; BF rise: p=0.58, t(6)=-0.58), implying its unique importance to SEA. Next, we applied a multinomial regression model to our data, using each of the elements (BF rise and PRN surge) to predict behavioral outcome in each trial. This model proved to be highly significant (χ^2 compared to the constant model: 17.51, p=1.6e-4; Supp. Table 1). Importantly, the PRN response was the strongest determining factor.

To examine the relationship between GRAB_NE_ and EEG/EMG dynamics above and beyond the effect on awakening, we calculated the correlation coefficients between response elements (BF rise and PRN surge), and EEG bands/EMG root mean square (RMS) for each mouse (Fig. 2J). Significant differences in correlation to the EEG sigma band (10-16 Hz) were found between PRN surge and BF rise (p=0.013, F(2)=5.58; p=0.01, post hoc PRN surge vs. BF rise). Correlations to EMG RMS were also significantly different, with PRN surge correlating positively and significantly above zero (p=4e-04, F(2)=12.39; p=4.4e-03, PRN surge vs. zero; p=5e-04, PRN surge vs. BF rise). These findings show that the PRN surge is uniquely anti correlated with the 10-16 Hz band and positively correlated with EMG amplitude, indicating a unique operational mode that goes beyond the binary classification of behavior.

To determine if the fast PRN NE activation stems from LC neuronal activity or if it is driven by dynamics at the target brainstem synapse, we recorded GCaMP7s from LC sub-populations projecting to the PRN. DBH-Cre mice were injected with AAVretro vectors encoding a Cre-dependent GCaMP7s in the PRN and BF (n_PRN_=7, n_BF_=7) and bulk calcium activity was recorded in the LC corresponding sub-population (Fig S3A). We found that the fast sound-evoked activity was already present to some extent in the activity of LC->PRN projecting neurons (Fig S3B), though that fast response was not indicative of trial outcome (data not shown). Thus, the result indicates that the PRN surge is partially explained by LC-> PRN neuronal activity.

### Early LC->PRN surge drives awakening

To determine if LC->BF and LC->PRN increased activity are sufficient to cause awakening and SEA we injected Cre-dependent AAVretro vector with ChR2, an excitatory opsin, to either BF (n=9) or PRN (n=9) and placed an optic fiber above the LC to induce specific activation of LC subpopulations (Fig 3A,B). First, we checked if optogenetic activation of each subpopulation promotes awakening (Fig 3C). In line with our results from GRAB_NE_ recordings, indicating similar activation in spontaneous awakening, in this experiment we found that both subpopulations are sufficient to induce awakening. We observed that activating the LC->PRN subpopulation induces awakening in a frequency-dependent manner (Fig 3E; p=5.85e-08, F(4)=16.18) with 40Hz stimulation being significantly more effective than lower stimulation frequencies (sham vs. 40Hz: p=2.8e-06; 5 vs. 40Hz: p=2.06e-07; 10 vs. 40 Hz: p=1.11e-06; 20 vs. 40 Hz: p=6.11e-04). LC->BF activation similarly depended on stimulation frequency (Fig 3D; p=0.004, F(4)=4.6,; sham vs. 40Hz: p=0.004, 5 vs. 40Hz: p=0.02, 10 vs. 40Hz: p=0.016).

**Figure 3:**
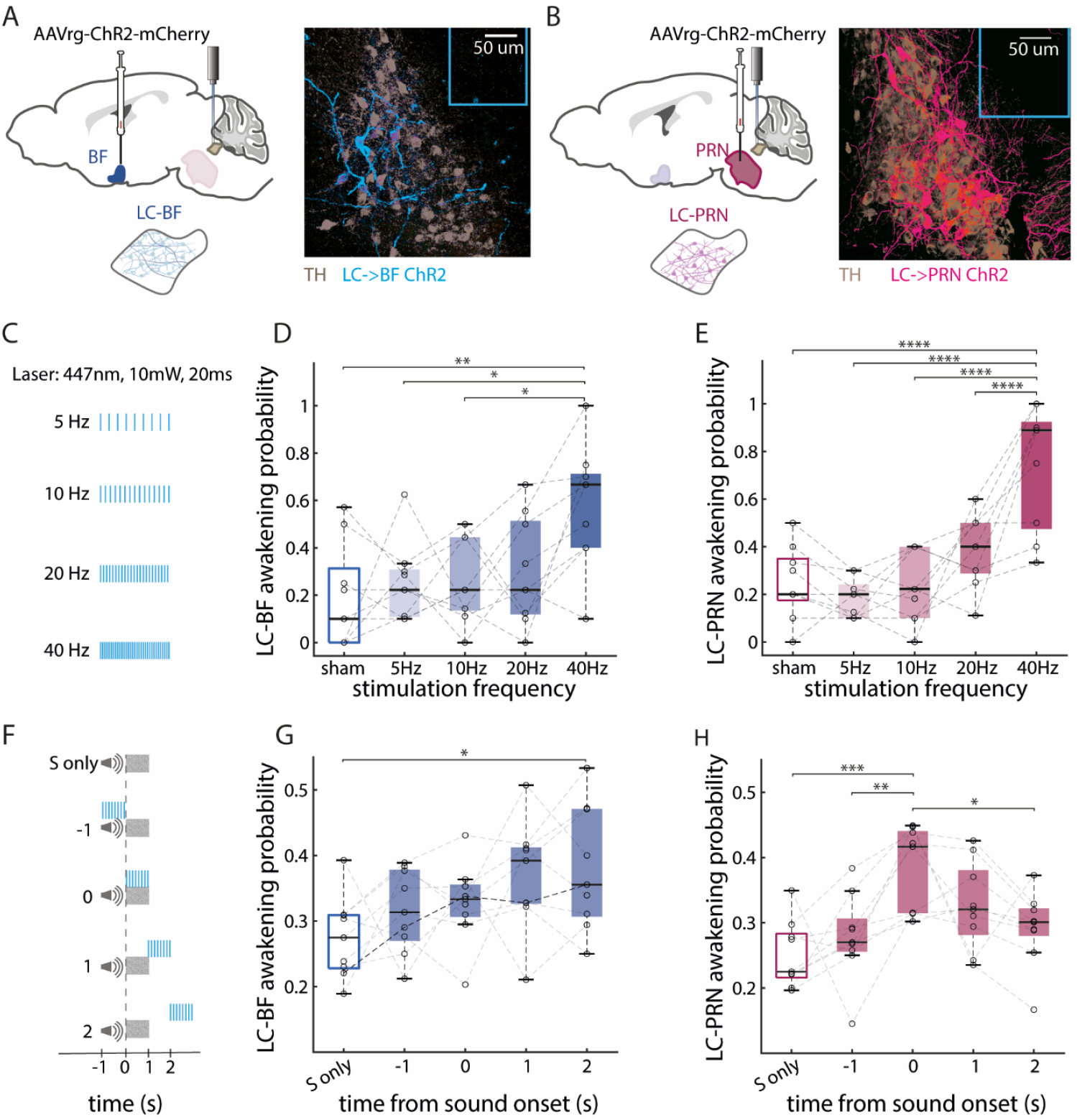
Early LC->PRN surge drives awakening. **A**, Depiction of surgical approach for LC->BF (left), and representative examples of fiber (cerulean line) and ChR expression in BF (right). Gray-TH, Blue-viral expression. **B**, Depiction of surgical approach for LC->PRN (left), and representative examples of fiber (cerulean line) and ChR expression in PRN (right). Gray-TH, Magenta-viral expression. **C**, Experimental procedure for laser awakening experiment. The experiment included 10s laser of 10mW and 20ms duty cycle for 5Hz, 10Hz, 20Hz and 40Hz. **D, E**, Box plot of probability to awaken from LC laser activation as a function of frequency in LC->BF ChR (**D**) and LC->PRN ChR (**E**). Dots represent single animals (n_BF_=9, n_PRN_=9). **F**, Experimental procedure for timing experiment. The experiment included either sound only (S only) or 1s laser of 10mW and 20ms duty cycle, 20Hz combined with a broad band noise in 80 dB SPL, so that the laser activation happened in different intervals relative to the sound-1s before, on time, 1s after or 2s after. **G, H**, Box plot of probability to awaken from LC laser activation as a function of timing in LC->BF ChR (**G**) and LC->PRN ChR (**H**). Dots represent single animals (n_BF_=9, n_PRN_=9).

Next, we investigated the effect of optogenetically activating LC->PRN and LC->BF during sleep when combined with sound presentation. Activation of 20 Hz occurred for 1s either before sound onset, together with sound presentation, or after sound presentation (Fig 3F). Given the distinct early peak in PRN NE and its relationship with SEA, we hypothesized that LC->PRN effects on awakening would depend on stimulation occurring together with sound, whereas LC->BF would be more strongly affected when stimulation occurs after sound offset. In line with this prediction, the effects of LC->BF optogenetic activation on awakening were associated with timing relative to auditory stimulation so that only stimulation of BF projecting LC neurons 2 secs after sound onset increased awakening probability (Fig 3G; p=0.029, F(4)=3.01; sound only vs. 2 secs: p=0.032). By contrast, LC->PRN activation showed a strong preference to laser stimulation that was synchronized with sound (Fig 3H; p=0.0009, F(4)=5.77; sound only vs. on time: p=0.0007, 1s before vs. on time: p=0.01, on time vs. 2s after: p=0.033: p=0.02).

Together with fluorophore-only control that show no effect (Fig S4), these results demonstrate that both subpopulations have a frequency and time dependency. The differences in the crucial timings of stimulation between the LC->BF and LC->PRN demonstrates the differences in these LC subpopulations. Furthermore, the timing of LC->PRN activation implies coherence and a more direct involvement in sensory processing, whereas the later LC->BF activation is more in line with the timing of the state changes than of the stimuli.

### Silencing the LC->PRN pathway strengthens EEG sigma power during NREM sleep

Finally, we set out to investigate whether NE release along specific projection pathways is necessary for effects on EEG and on awakening probability. To this end, we selectively silenced synaptic release of LC neurons in either BF or PRN target regions using the inhibitory OptoGPCR PdCO^38^ (n=8 DBH-Cre mice). We injected a Cre-dependent PdCO viral construct to the LC and placed optic fibers above both the PRN and BF target regions (Fig 4A-C). We first examined the effect of silencing on otherwise uninterrupted sleep to analyze spontaneous awakening. Silencing LC synapses simultaneously in both areas consolidated NREM sleep, as evident by lower probability to transition to wakefulness (Fig 4D,E; p=0.05, F(3)=2.94; p=0.05, sham vs. both). Furthermore, silencing LC->PRN synapses alone during NREM sleep significantly elevated EEG power in the low frequencies of the sigma band (9.5 - 12.25Hz), while silencing both PRN and BF pathways simultaneously additionally led to a power increase in the higher end of the sigma band (14.2-16.8 Hz), together with increase in the power of 20.7-22.35 Hz (Fig 4F). This elevation in power, specifically in the sigma band, points to a more consolidated and resilient sleep^39,40^. A comparison of average power in the combined cluster confirmed this conclusion (Fig 4G; p=2e-04, F(3)=9.16; sham vs. BF: p=0.06; sham vs. PRN: p=0.003; sham vs. both: p=2e-04).

**Figure 4:**
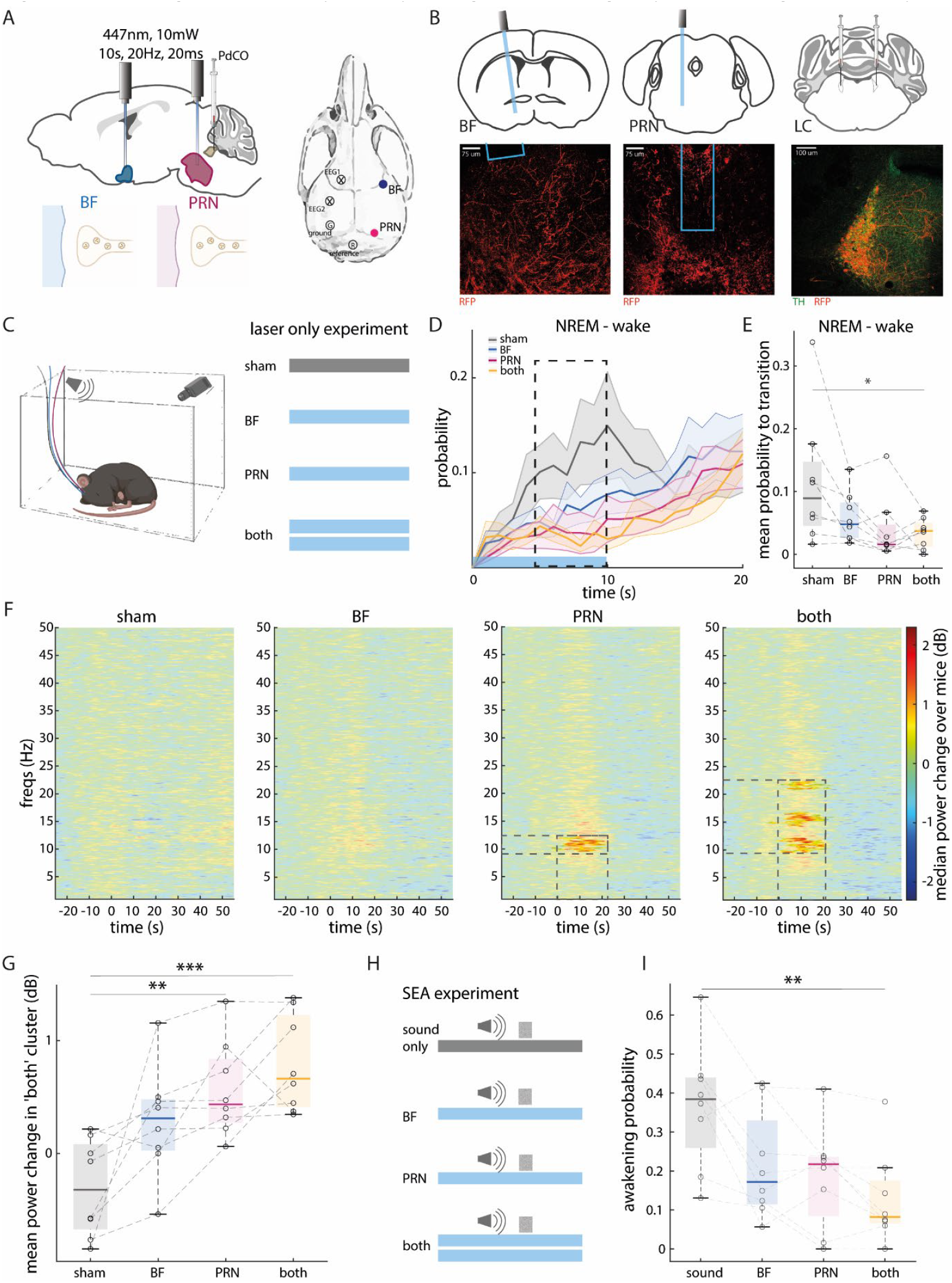
Silencing the LC->PRN pathway strengthens EEG sigma power during NREM sleep. **A**, Depiction of surgical approach. **B**, Representative examples of fiber and PdCO expression (RFP) in BF (left), PRN (middle) and LC. In BF and PRN where the fibers were placed the expression shows LC axons in red. In the LC the expression is seen in the cell bodies: Green-TH, red-viral expression. **C**, Left-experimental setup, right-experimental procedure. The experiment included 10s laser of 10mW and 20ms duty cycle 20Hz stimulation activated in BF, PRN, both simultaneously and sham trials. Animals were freely moving in their home cage at the time. **D**, Probability to transition from NREM to wake given that in t=0 the state was NREM. Gray-sham, blue-BF, magenta-PRN, yellow-both. Laser was on from t=0 to t=10s, depicted by cerulean line in bottom. dashed square represents the time taken for further analysis (t=5-10s). **E**, Box plot of average probability to transition from NREM to wake under each condition during t=5-10 from laser onset as marked in panel D. **F**, Median EEG spectrograms across mice around trials during NREM normalized to t=-30-0s relative to laser onset. Laser was on from t=0 to t=10s. left-sham condition, BF laser condition, PRN laser condition, right-both lasers on condition. Areas that are not significant compared to sham (see methods) are opaque. Areas that are significant are fully colored and have dashed squares around them. **G**, Average mean power change within the significant cluster found in “both” in panel F in each of the conditions. Gray-sham, blue-BF, magenta-PRN, yellow-both. **H**, Experimental procedure. The experiment included 10s laser of 10mW and 20ms duty cycle 20Hz stimulation activated in BF, PRN, both simultaneously and sham trials. At t=8s broad band noise of 80 dB SPL was played by a mounted speaker for 1s. **I**, Box plot of awakening probability.

Next, we examined the effect of silencing LC synapses in target regions with respect to sound evoked awakening (-8 +2s around sound onset, Fig 4H). Such silencing lowered the probability to awaken by external stimulus (Fig 4I, p=0.012, F(3)=4.4). Evoked awakening probability, when simultaneously silencing both target regions, was significantly lower than sham (p=0.008, Tuckey Kramer post hoc test). PRN silencing showed a trend (p=0.06, Tuckey Kramer post hoc test) for reduced awakening probability, while silencing the BF alone was not significantly different compared to sham stimulation (p=0.14, Tuckey Kramer post hoc test). Fluorophore-only control experiments confirmed that laser application itself was not associated with significant changes in awakening probability (Fig S5).

Together, these projection-specific silencing experiments show that silencing LC synaptic release in both the PRN and BF impacts the EEG and reduces awakening probability. Effects are strongest for combined silencing in the two targets, while specifically silencing PRN is associated with stronger effects.

## Discussion

We investigated whether neuronal and behavioral responses to sounds during sleep recruit the LC-NE system globally or via projection-specific pathways. To this end, we examined arousal-promoting LC-NE pathways involving the brainstem (PRN) and the basal forebrain (BF). Our results establish that these LC-NE pathways differentially contribute to sensory-evoked awakening (SEA) during sleep, with an early surge of activity in the LC->PRN pathway playing a privileged role. We show that LC-NE heterogeneity arises both in terms of the LC neurons (when monitoring or manipulating specific subpopulations via AAV retro vectors), as well as through NE dynamics in target regions (when monitoring extracellular NE using GRAB_NE,_ or manipulating synaptic release via PdCO). Together, the process of sound-evoked awakening is predominantly modulated by an early activity of LC->PRN projecting neurons and NE release and dynamics at the brainstem, attesting to modularity in LC-NE signaling even in the context of sleep and arousal.

Anatomically, LC neurons targeting the forebrain are located more dorsally compared to those projecting to the PRN (Fig 1). This dorsal-to-ventral gradient is reminiscent of past anatomical studies that indicate that the LC is organized by projection target^23,41^. Anatomical distinction was further reflected physiologically (Fig 2), with sound-evoked NE activity patterns differing significantly between the PRN and the BF. Specifically, NE profiles at the PRN exhibited a strong rapid surge of activity that was tightly synchronized with the sensory stimulus, and predicted whether the animal would awaken or remain asleep within seconds. Optogenetic manipulation (Fig 3) established the causal role of rapid NE surges in the LC->PRN pathway during SEA. Inducing this surge concurrently with auditory stimuli significantly increased awakening probability. Moreover, silencing LC->PRN synaptic release (Fig 4) exhibited some signs of consolidating NREM sleep to a greater extent than LC->BF silencing, further supporting the notion that the LC->PRN pathway is important in promoting awakening.

These findings highlight multi-layered heterogeneity within the LC-NE system, promoting the view that arousal signaling is circuit-specific rather than globally uniform. Accordingly, LC-NE modularity is evident both in NE dynamics at target regions (as evident when using GRAB_NE_ and PdCO) and in activities of presynaptic LC sub-populations based on their projections (as evident using AAVretro vectors with GCaMP7s and ChR2). Thus, heterogeneity in the LC-NE system requires investigation of several complementary mechanisms occurring simultaneously pre-synaptically, at the synapse, at post-synaptic receptors, and at non-neuronal compartments^17,42,43^. Our results join previous work that showed LC modularity in the context of other functions such as anxiety, pain, and learning circuits^21,34^, and extend it for the first time to demonstrate such heterogeneity in the context of arousal and wake-promoting circuits.

What underlies the specific fast surge of NE activity in the PRN? For long, a distinction has been made between tonic (slow) and phasic (fast) LC-NE activity patterns^44^. Tonic firing is more related to slow dynamics internal states and arousal, such as vigilance states^45^ and dynamics in the order of seconds or tens of seconds occurring during wakefulness^46,47^ and NREM sleep rhythm^7,48^. Phasic firing is more related to sensory responses, decision-making, and network switching^12,44,49,50^. Indeed, a recent whole-brain optogenetics-fMRI study in mice revealed that LC surges are more likely to recruit regions involved in sensory processing^43^. It is therefore likely that the PRN NE surge we observe following sound presentation during sleep represents phasic LC firing in response to salient surprising stimuli. Phasic activity may express more strongly in adjacent brainstem regions as unmyelinated LC axons serve as a low-pass filter for distant NE signaling. Finally, the findings indicate that SEAs are more effective in revealing LC-NE heterogeneity than spontaneous awakenings. This could possibly be related to sensory-evoked phasic activity rather than changes in tonic activity levels around spontaneous state transitions. Indeed, our results indicate that while both pathways reflect tonic-like state dependent changes, the PRN is unique in displaying the phasic-like sensory evoked response.

A recent study showed that the PRN is involved in an auditory pathway implicated in SEA^51^. Indeed, the PRN receives input from the cochlear nucleus and has downstream targets involved in auditory responses during both NREM sleep and wakefulness^52^. Our results further support this notion and highlight that, in addition to ascending auditory signaling via glutamatergic pathways, NE modulation at the PRN supports its key role in integrating arousal information into sensory pathways. Future research could test whether LC-NE PRN activity also generalizes to arousal and awakening signaling in other sensory modalities beyond auditory.

Some limitations in the current study could be overcome in future studies employing refinements in research methodologies. At present, our silencing of projection-specific LC pathways with PdCO only provides temporal resolutions of many seconds. Thus, it was not possible to precisely time this intervention together with short auditory stimulation. Future studies could harness projection-specific silencing tools with higher temporal precision to test the effects of silencing LC->PRN at the specific windows of the early 0-1 sec surge. Additionally, bilateral silencing in PRN target regions is challenging due to the proximity of optic fibers on the mouse brain. Future studies could explore such bilateral interventions, which have potential of eliciting stronger effects. Our investigation focused on the auditory modality, building on previous work associating sound-evoked awakening with LC-NE activity^14^. Future studies could also test sensory-evoked awakening along other modalities (e.g., somatosensation, olfaction), and seek to generalize findings to females. Finally, the current study focused on the PRN and the BF as representing target regions in the brainstem, and forebrain, respectively. Future work can explore additional LC projection pathways known to be involved in arousal, such as the thalamus^8,53^ and the hypothalamus^54^.

In conclusion, our findings reveal that an early surge of NE signaling in the LC->PRN pathway plays a pivotal and specialized role in triggering awakening in response to sounds. More generally, this finding highlights the specific role that LC projection pathways may have in mediating arousal and other functions. This research underscores the modular nature of LC-NE functions and paves the way for further exploration of the specific roles of downstream arousal targets.

## Resource availability

### Lead contact

Requests for further information and resources should be directed to and will be fulfilled by the lead contact, Noa Matosevich.

Any additional information required to reanalyze the data reported in this paper is available from the lead contact upon request.

## Acknowledgements

We thank all our laboratory members as well as Prof. Inna Slutsky, Prof. Ofer Yizhar, Dr. Yaniv Sela and Dr. Hanna Hayat for thoughtful discussions and comments on the manuscript. We thank Flavio Schmidig for his assistance in developing the EEG analysis code.

This work is supported by grants from the Israel Science Foundation (ISF 1557/22) and the European Research Council (ERC-2019-COG 864353).

## Author contributions

Conceptualization, N.M., N.R. and Y.N.; Methodology, N.M. and N.R.; Software, N.M. and A.M.; Formal Analysis, N.M., S.K. and A.M.; Investigation, N.M., U.K., N.Z., S.K. and N.G.; Data Curation, N.M., U.K., N.Z., S.K. and N.G.; Writing-Original Draft, N.M.; Writing-Review & Editing, Y.N., N.R., N.Z. and U.K.; Visualization, N.M., Supervision, N.R. and Y.N.

## STAR Methods

### Experimental Model

#### Animals

All experimental procedures including animal handling, surgery, and experiments followed the NIH Guide for the Care and Use of Laboratory Animals and were approved by the Institutional Animal Care and Use Committee (IACUC) of Tel Aviv University (approval 01-19-037, 01-22-004). Adult c57bl/6j (RRID:MGI:3028467) mice (8-12 weeks old at the time of surgery) were used for retrobeads and GRAB_NE_ experiments. Adult DBH-cre mice (B6.FVB(Cg)-Tg(Dbh-cre)KH212Gsat/Mmucd, RRID:MMRRC_036778-UCD; 036778-UCD-HEMI) were used for all other experiments. For viral tracing, four mice were female, and one was male; all other experiments were performed on male mice. Mice were housed in transparent Plexiglas cages at constant temperature (20°-23°C), humidity (40-70%) and circadian cycle (12h light/dark cycle, starting at 10:00 AM). Food and water were available ad libitum.

### Method details

#### Surgery

The coordinates used to target BF were: (respective to bregma) anteroposterior (AP): 0.7 mm; medio-lateral (ML): 2.25 mm at 7.5°, dorso-ventral (DV): -5 mm relative to the brain surface. The coordinates used to target PRN: (respective to bregma) anteroposterior (AP): -4.25 mm; medio-lateral (ML): 1 mm; dorso-ventral (DV): -4.8 mm relative to brain surface. Coordinates used to target the LC: (respective to bregma) anteroposterior (AP): - 5.4 mm; medio-lateral (ML): 1 mm; dorso-ventral (DV): -3.15 mm relative to brain surface.

For anatomical tracing, Red Fluorescent Latex Microspheres (retrobeads; Lumafluor) were used as a general retrograde tracker in a total volume of 300 nl injected into PRN or BF. The Cre-dependent retro viruses were used as a specific noradrenergic retrograde neuronal tracer and injected into both regions-EGFP to the PRN and mCherry to the BF in a total volume of 500 nl for each injection (Fig 1A,E). For simultaneous GRAB_NE_ experiments, after injection of GRAB_NE_ virus to PRN and BF, optic fibers (MFC_400/430-0.48_5mm_MF1.25_FLT, doric) were lowered to the same coordinates 0.15mm higher in the DV axis (Fig 2A). For retro ChR2 experiments, a cre-dependent retro AAV with ChR2 was injected to either the PRN or the BF (as described above), and an optic fiber placed above the LC (MFC_200/240-0.22_5mm_MF1.25_FLT, doric; Fig 3A,B). For PdCO experiments, the LC was injected with a cre-dependent AAV with PdCO or fluorophore only (for control experiment, Fig S5) as described above. 3-5 weeks after injection optic fibers (MFC_200/240-0.22_5mm_MF1.25_FLT, doric) were implanted above PRN and BF (Fig 4A).

#### Experiments

At least one week of habituation was allowed between surgeries and experiments after which mice were placed in a home cage within a sound-attenuation chamber (-55dB, H.N.A) and connected to the EEG/EMG headstage and to the fiberoptic patch cord (MFP_400/430/1100-0.57_1m_FC-ZF1.25_LAF, doric lenses) for photometry recordings, and (MFP_200/240/900-0.22_1m_FC-MF1.25, doric lenses) for optogenetic manipulations. The camera and speaker were mounted 50 cm above the cage floor (Fig 2A). After > 48h in the new cage, habituation also gradually introduced tethering and exposure to sounds. <u>Optogenetics awakenings</u>(Fig S1E-F) -Laser was triggered manually by the experimenter when the mouse was in NREM sleep (as observed using EEG, EMG and video feed). Laser parameters were set to 10 s duration, 10 Hz frequency and 90 ms duty cycle. <u>Undisturbed sleep-</u> (Fig S2, Fig 2C) Electrophysiology and photometry data were recorded continuously for 12 hours (starting around light onset 10AM) while the mice were behaving freely. <u>Sound evoked arousal (SEA</u>)-(Fig 2D-J, Fig 3F-H, Fig 4H-I, Fig 34, Fig S5C) BBNs as described above were presented intermittently in randomized order every 60 secs (±0.5s jitter). <u>Laser awakening experiment</u> – (Fig 3C-E, Fig S4) laser parameters: 10s, frequency: 1,5,10,20 and 40 Hz. Each parameter ran 10 times. The order of trials was randomized. Trial start was initiated manually so that all trials occurred during NREM after at least 15 secs of continuous NREM with at least 30 s between trials. <u>Timing SEA experiment</u>– (Fig 3F-H) To determine the criticality of synchronization between LC subpopulation activity and auditory stimulation we changed the laser activation timing relative to sound. laser parameters: 1s ,20 Hz. Sound parameters: broadband noise (BBN) bursts (1sec duration 80 dB SPL). Every 60sec (± 0.5sec jitter) a trial was randomly initiated in which either laser started 1 s before sound, in time with sound, 1 s after sound or 2 s after sound onset. <u>PdCO experiments</u>- (Fig 4) 447 nm laser was set to 10 mW at fiber tip, 10s stimulation, 20 Hz, 20ms. <u>Laser only experiment</u>- (Fig 4C-G) After habituation, mice were recorded, and stimulation was applied every 2 minutes (with 0.5 s jitter). Stimulation was either in the BF, the PRN or both. Sham onsets were randomly selected from the data so that there were the same number of trials as in both site stimulation, and the times were between 50 and 70 seconds from other trial onsets. <u>SEA + laser experiments-</u> (Fig 4H-I) Stimulation was applied every 2 minutes (with 0.5 s jitter). Stimulation was either in the BF, the PRN, both or none. At t=8s from laser onset 1s BBN was played (80 dB SPL).

#### Histology

Viral expression or retrobead localization was evaluated histologically by examination of double staining of free-floating sections. To this end, primary antibodies were against tyrosine hydroxylase to localize LC cells or RFP to improve PdCO viral expression visibility in axons (chicken anti-TH 1:300, ab76442 abcam, RRID: AB_1524535; guinea pig anti RFP 1:500, 390004 SYSY, RRID:AB_2737052), and secondary antibodies conjugated to fluorophores were in appropriate coloring so as not to overlap with the viral fluorophore (AF405 goat anti-chicken 1:200, ab175674 abcam, RRID:AB_2890171; AF488 goat anti-chicken 1:500, ab150173 Abcam, RRID:AB_2827653; AF488 goat anti-chicken 1:50, A11039 Invitrogen, RRID:AB_2534096; donkey anti guinea pig Cy^TM^5 1:300, 706-175-148 /Jackson ImmunoResearch, RRID:AB_2340462). Images were acquired by a LEICA SP8 high-resolution laser scanning confocal microscope (Leica, Wetzlar, Germany) and a x10 air/0.4 NA objective, and contrast and brightness were improved for representative images.

### Quantification and statistical analysis

Unless stated otherwise, statistical analysis was done using ANOVA and Tuckey Kramer post hoc analysis. Shaded areas in graphs represent SEM. In box plots, the central mark in each box indicates the median, and the bottom and top edges of the box indicate the 25th and 75th percentiles, respectively. The whiskers extend to the most extreme data points not considered outliers. Dashed lines represent single animals. *p<0.05, **p<0.01, ***p<0.001, ****p<1e-04. illustrations were created using BioRender and Adobe Illustrator.

#### Fiber photometry

Data were detrended and normalized in periods when LED was active as described in^7^. In short, the isosbestic channel was fitted to the GRAB trace using a linear polynomial fit, then used as the f_0_ reference point: 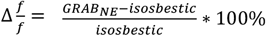 (Fig S2).

#### Sleep scoring

Scoring of sound evoked awakenings (SEA) was performed offline while visualizing EEG, EMG, and video data in TDT ‘scope’ software so that scorer was blinded to trial intensity. Each trial was given two scores: one characterizing the state at trial onset (wake/NREM/REM), and the second categorized the behavioral outcome (maintained/awakening/EEG activation/EMG activation/short awakening). Baseline states were determined based on the state of the animal at the 5 seconds preceding trial. Cases in which the state was not stable at that time window were discarded. Behavioral categorization was determined based on the response from trial onset until 3 seconds later. Maintained sleep was declared only if there was no visible change in EEG, EMG and video (38.14% of trials). Awakening was declared only in the case of EEG activation, EMG activation and movement observed in the video that lasted at least 3 seconds (32.72% of trials; Fig 2D). If all 3 conditions were observed for a period shorter than 3 secs, the trial was tagged as ‘short awakening’ (14.92% of trials). If only one was met the trials were tagged as ‘EEG activation’ (5.63%;) or ‘EMG activation’ (8.59%) in accordance with the activated channel. Given the relatively lower probabilities and inconsistency of ‘semi-awakening’ trials (e.g., short, EEG and EMG activation), those were eventually left out of analyses.

Undisturbed or laser-only recording sessions (without auditory stimulation) were scored continuously using semi-automatic sleep scoring as in^55^ (adapted from^56^). A 1h+ segment of manually scored data in each session was used in an automated sleep-scoring algorithm as in using a convolutional neural network (CNN) that was trained using >20 manually scored blocks from prior mice recordings in the lab. The algorithm was fed with the manual sleep scoring vector (tagged in 1s resolution) along with the parietal EEG signal (low-pass filtered < 20Hz), and the EMG signal (band-pass filtered 10Hz – 50Hz). The entire data set was classified into sleep scoring labels (wake/NREM/REM). Automatic classification of vigilance state was visually inspected to ensure accuracy (Fig S2E).

#### Analysis of anatomical profiles of LC->PRN and LC->BF subpopulations

##### Anatomical gradients of LC->PRN vs. LC->BF subpopulations based on red fluorescent Retrobeads data (Fig 1A-D)

For each animal, every coronal slice was localized to coordinates relative to Bregma according to the Allan Brain Atlas. In every image, the LC dimensions based on TH immunostaining were manually marked. The image was divided into DV strips of 100 µm. Cells within the each DV strip in every coronal slice LC were manually counted. Since every animal had different available coronal slices and the number of these available slices were varied, each cell count was weighted by the number of animals that contributed to it. Furthermore, to normalize the number of cells per animal, the count was divided by the average count for the same locations as well as the ratio between the animal yield and the expected yield. Then the normalized values were multiplied by the average cell count to establish the cell proportion values. To determine statistical significance, a Monte-Carlo permutation test was employed. First, the absolute difference between the two subpopulations was calculated as the square root of the mean squared difference between the PRN and the BF distributions. Then the same measurement was calculated on 100,000 randomly assigned permutations. The number of animals in each group was fixed, but the allocation of animals to every group was randomized. This created a distribution of possible mean absolute differences and allowed us to calculate the one-tailed p value associated under the hypothesis that the distributions are different. Upon determining a significant difference, we ran the same test to discover if the difference was in the DV axis or the AP axis, or some combination. For each population, the mean location in each axis was calculated and the difference between PRN and BF was used as the value to be compared. The resulting two-tailed p value was calculated from the distribution.

##### Co-localization of LC->PRN vs. LC->BF neurons based on retrograde virus tracing (Fig 1E-G)

The double-labeling viral tracing quantification of the extent of colocalization of the red and green channels was carried out using Imaris Software (Imaris 9.0.1, RRID:SCR_007370). For each coronal slice, the LC boundaries were defined based on the TH immunostaining. Cells in the two colors (red and green) within the LC were selected and were automatically counted. The number of co-localized cells was identified using the ‘Colocalize spots’ tool (with a maximum distance of 10 um between the red and green spots). The position of each slice at the anterior-posterior axis, relative to the Bregma, was manually matched to its corresponding position in the Allen Mouse Brain Atlas. For each slice, the ratio of colocalized cells to the total number of cells was calculated for both the BF and PRN projecting cells. The mean ratio of PRN and of BF colocalized cells per mouse was then obtained.

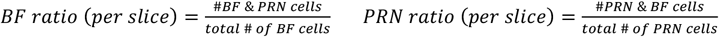

To determine statistical significance, two two-sample t-tests were applied to test whether each population is a sub-sample of the other. Thus, the mean ratio of colocalized cells per mouse for each subpopulation was tested against 1 (1 meaning all BF cells are also PRN cells or vice versa).

#### Analysis of simultaneous GRAB_NE_ data in PRN and BF, and of retro-GCaMP

We examined average traces (across trials) of PRN and BF GRAB_NE_ after 5s baseline subtraction (Fig 2C,E-J, Fig S3B). To confirm that the baselines did not hold significant information we compared the baselines under each outcome across mice using ANOVA1 in PRN (f(3)=0.15, p=0.93) and BF (f(3)=0.62, p=0.6). To determine temporal intervals associated with significant differences between NE dynamics in PRN and BF (horizontal bars at the bottom of Fig. 2f), we divided data to 0.3 sec bins and compared the resulting 13 bins between t=0 and t=4 s separately. Upon getting a significant Kruskal-Wallis test result (BF: *χχ*^2^(13) = 55.6, p=3.17 e-07; PRN: *χχ*^2^(13) = 41.32, p=8.45 e-05) we ran a one-sided multiple comparisons correction using Dunnetts’ test to determine which bins were significantly above 0. Consecutive significant bins were grouped together, and the result was two significant time bins in the PRN trace: 0.3 - 0.6 s (‘PRN surge’) and 3 - 3.9 s (‘PRN rise’); and 1 significant time bin in the BF trace: 1.8 - 3.9 s (‘BF rise’). <u>Data for spontaneous awakening</u> from NREM sleep (Fig. 2C) were obtained from sessions in which no sounds were played, scored continuously (details above). Awakening trials were normalized to their -15: -10 s baseline.

#### EEG power analysis

EEG power analysis was performed on the parietal EEG channel using the ‘newtimef’ function in MATLAB. Mean event-related (log) spectral perturbation (ERSP) was calculated on the trial matrix of -30 – 60 s around laser onset in each state (wake/NREM/REM) using FFTs and Hanning window tapering. The maximum window size is 10 s. The frequencies calculated from 1 to 50 Hz. Single trial normalization was not applied (fig 4f, fig 2J). The Median ERSP over mice in each laser condition was calculated. Representational dissimilarity analysis was performed using the FieldTrip toolbox^57^. In short, using one EEG channel, t-tests were run comparing each condition with the sham condition. Final statistic was calculated using a Monte-Carlo permutation test with cluster correction. Cluster alpha and configuration alpha were set to 0.1 for a one-sided test with a parametric cluster threshold. The number of randomizations was set to 1000.

#### State transition analysis

Awakening probability was calculated as the probability of full awakening over all NREM trials under the specific parameter (fig 3E, F, H, I, fig 4J). Statistics were carried out using anova1 with post hoc Tuckey Kramer tests. For every PdCO mouse, the state it was in in each trial time point was added and divided by the number of trials that had the same base state at 5 s before trials onset. This resulted in a data structure containing the probability of each state from trial onset to 60 s after under the base state condition (fig 4D). Data was divided into dark and light phases and after observing no discernable differences only light phases results are displayed.

### Key resources table

| REAGENT or RESOURCE | SOURCE | IDENTIFIER |
| --- | --- | --- |
| **Antibodies** |  |  |
| chicken anti-TH 1:300, ab76442 | abcam | RRID: AB_1524535 |
| guinea pig anti RFP 1:500, 390004 | SYSY | RRID: AB_2737052 |
| AF405 goat anti-chicken 1:200, ab175674 | abcam | RRID: AB_2890171 |
| AF488 goat anti-chicken 1:500, ab150173 | abcam | RRID: AB_2827653 |
| AF488 goat anti-chicken 1:50, A11039 | Invitrogen | RRID: AB_2534096 |
| donkey anti guinea pig Cy<sup>TM</sup>5 1:300, 706-175-148 | Jackson ImmunoResearch | RRID: AB_2340462 |
| **Virus strains – viruses were produced in Viral Vector Facility (VVF) of the Neuroscience Center Zurich (ZNZ)** |  |  |
| ssAAV-retro/2-hSyn1-dlox-EGFP(rev)-dlox-WPRE-hGHp(A) | pAAV-hSyn-DIO-EGFP was a gift from Bryan Roth | Addgene viral prep # 50457-AAVrg; RRID: Addgene_50457 |
| ssAAV-retro/2-hSyn1-dlox-mCherry(rev)-dlox-WPRE-hGHp(A) | pAAV-hSyn-DIO-mCherry was a gift from Bryan Roth | Addgene viral prep # 50459-AAVrg; RRID: Addgene_50459 |
| AAV2-hSyn1-GRAB(NE1m)-WRPE-hGHp(A) | pAAV-hSyn-GRAB_NE1m was a gift from Yulong Li<sup>36</sup> | Addgene viral prep # 123308-AAV9; RRID: Addgene_123308 |
| pAAV-hSyn-GRAB_NEmut | pAAV-hSyn-GRAB_NEmut was a gift from Yulong Li<sup>36</sup> | Addgene viral prep # 123310-AAV9; RRID: Addgene_123310 |
| CAV2-PRS-ChR2-mCherry | Eric Kremer | N/A |
| ssAAV-retro/2-hEF1α-dlox-hChR2(H134R)_mCherry(rev)-dlox-WPRE-hGHp(A) | pAAV-EF1a-double floxed-hChR2(H134R)-mCherry-WPRE-HGHpA was a gift from Karl Deisseroth | Addgene viral prep # 20297-AAVrg; RRID: Addgene_20297 |
| ssAAV-1/2-hSyn1-lox71-ePdCO_mScarlet(rev)-lox66-WPRE-hGHp(A) | pAAV-hSyn-SIO-PdCO-mScarlet-WPRE was a gift from Ofer Yizhar<sup>38</sup> | Addgene plasmid # 198509; RRID: Addgene_198509 |
| ssAAV-5-hSyn1-dlox-mCherry(rev)-dlox-WPRE-hGHp(A) | pAAV-hSyn-DIO-mCherry was a gift from Bryan Roth | Addgene viral prep # 50459-AAV5; RRID: Addgene_50459 |
| **Experimental models: Organisms/strains** |  |  |
| Mouse: B6.FVB(Cg)-Tg(Dbh-cre)KH212Gsat/Mmucd; 036778-UCD-HEMI | The Jackson Laboratory | RRID: MMRRC_036778-UCD |
| Mouse: C57BL/6J | The Jackson Laboratory | RRID: MGI:3028467 |
| **Software and algorithms** |  |  |
| MATLAB | Matlab Mathworks | RRID: SCR_001622 |
| Imaris 9.0.1 | Imaris | RRID: SCR_007370 |
| Synapse | Tucker-Davis Technologies | RRID: SCR_006495 |
| Scope | Tucker-Davis Technologies | RRID: SCR_006495 |
| Biological animations | BioRender | RRID: SCR_018361 |
| **Other** |  |  |
| Red Fluorescent Latex Microspheres (retrobeads) | Lumafluor | N/A |
| 1-site Fiber Photometry system | Doric Lenses, Canada | N/A |
| 4-ports minicube | Doric Lenses, Canada | FMC4 or iFMC4 |
| Femtowatt photoreceiver | Newport | 2151 |
| amplifier | Tucker-Davis Technologies | PZ5 |
| BioAmp Processor | Tucker-Davis Technologies | RZ2 |
| Auditory stimuli generation and amplification | Tucker-Davis Technologies | SA1 |
| magnetic speaker | Tucker-Davis Technologies | MF1 |

