## supplementary methods for "An early surge of norepinephrine along brainstem pathways drives sensory-evoked awakening"

### Animals

The mouse strain used for this research project, B6.FVB(Cg)-Tg(Dbh-cre)KH212Gsat/Mmucd, RRID:MMRRC_036778-UCD, was obtained from the Mutant Mouse Resource and Research Center (MMRRC) at University of California at Davis, an NIH-funded strain repository, and was donated to the MMRRC by MMRRC at University of California, Davis. Made from the original strain (MMRRC:032081) donated by Nathaniel Heintz, Ph.D., The Rockefeller University, GENSAT and Charles Gerfen, Ph.D., National Institutes of Health, National Institute of Mental Health[58,59]

### Method details

#### *Surgery*

Before all surgical procedures, mice were anesthetized (with isoflurane 3% by volume for induction and 1.3% for maintenance) and placed in a stereotaxic frame. Surgery was performed in aseptic conditions and the mice received antibiotics (Cefazoline 15 mg/kg s.c) and Carprofen/Rimadyl 5 mg/kg i.p. For virus injections and optic fiber implantations, a craniotomy was performed under microscopic control using a high-speed surgical drill. Injections were made using a UMP3 Microsyringe Injector and a Micro4 Controller pump, and 33-gauge NanoFil needles World Precision Instruments, USA at a 100 nl per minute pace. At the end of the injection the scalp was sutured and the mouse returned to his home cage for 48-72 hours to allow for retrobead tracing, 3 weeks for retro viral tracing.

At the end of all optic fiber implantation surgeries, dental acrylic was gently placed around the optic fibers, fixing them to the skull. Two screws (one frontal and one parietal; 1mm in diameter), were placed over the left hemisphere for the EEG recording, and two additional screws were placed above the cerebellum and posterior parietal lobe as reference and ground (Fig 4A). EMG was measured via two single-stranded stainless-steel wires inserted to either side of the neck muscles in a bipolar reference configuration. EEG and EMG wires were soldered onto a custom-made omnetics headstage connector. Metabond dental cement was used to cover all screws and EEG/EMG wires.

#### *EEG, EMG, and video recordings*

EEG and EMG were digitally sampled at 1,017 Hz (PZ5 amplifier, Tucker-Davis Technologies) and filtered online: Both EEG and EMG signals were notch-filtered at 50Hz and 100Hz to remove line noise and harmonics; then, the EEG and EMG signals were band-pass filtered at 0.5-200Hz and 10-100Hz, respectively. Simultaneous video data (for sleep and basic behavioral assessments) during free behavior were captured by a USB webcam synchronized with electrophysiology/photometry data.

#### *Fiber photometry*

Fiber photometry data was collected as described in [60]. Briefly, using a 1-site Fiber Photometry system (Doric Lenses, Canada) adapted to two excitation LEDs at 465nm (GCaMP, $GRAB_{NE}$) and 405nm (isosbestic control channel). Simultaneous monitoring of the two channels was made possible by connecting the LEDs to a 4-ports minicube (with dichroic mirrors and cleanup filters to match the excitation and emission spectra; FMC4 or iFMC4, Doric Lenses) via an attenuating patch cord (400 µm core, NA=0.37-0.48). LEDs were controlled by drivers that sinusoidally modulated 465nm/405nm excitation at 217/330 Hz, respectively, enabling lock-in demodulation of the signal (Doric Lenses, Canada). Zirconia sleeves were used to attach the fiber-optic patch cord to the animal's cannula. Data were collected using Femtowatt photoreceiver 2151 (Newport) or through an integrated photodetector in the case of iFMC4 and demodulated and processed using an RZ2 BioAmp Processor unit and Synapse software (Tucker-Davis Technologies). The signal, originally sampled at 24,414Hz, was demodulated online by the lock-in amplifier implemented in the processor, sampled at 1,017.25Hz and low-pass filtered with a corner frequency at 6Hz. All signals were collected using Synapse software (Tucker-Davis Technologies). To achieve 12/24-hour recordings with minimal adverse effects of prolonged LED activation (e.g., phototoxicity and photobleaching), we automatically turned the LEDs off every hour and allowed an hour with LEDs off (Fig S2).

#### *Laser parameters for optogenetic experiments*

Blue stimulation at 447 nm for optogenetic excitation and silencing was delivered via lasers (CNI, China) coupled to optic fibers whose timing and intensity were automatically controlled via RZ2 (TDT). Light intensity at optic fiber tips was measured with a power meter (ThorLabs PM100D) before optic fiber insertion and set to 10mW.

#### *Auditory stimulation parameters*

Sounds were generated in TDT software and amplified (SA1, Tucker Davis Technologies (TDT)), and played free field through a magnetic speaker (MF1, TDT). Broadband noise (BBN) bursts of 1sec duration (0.8 V peak to peak waveform), either 73, 80, or 88 dB SPL, order randomized (27.94 ± 3.6%, 34.44 ± 4.34% and 38.16 ± 4.26% awakening respectively) were presented intermittently (± 0.5sec jitter). Sound intensities were measured by placing a Velleman DVM805 Mini Sound Level Meter at the center of the cage floor.

#### *Histology*

Following all experiments, under deep isoflurane anesthesia (4%) combined with a ketamine-xylazine dose (ketamine 100 mg/kg, xylazine 1.33 mg/kg), the mice were perfused intracardially with saline (0.9% NaCl; 1 ml/g) followed by 4% paraformaldehyde (Merck). Their brains were then extracted and fixed for 24 - 48 h in 4% PFA. Coronal brain sections were cut using a Leica VT1000 S vibrating blade microtome at 60µm. Then, sections were either kept free floating in PBS for immunohistochemistry (IHC) or mounted on glass slides and examined under bright field microscopy.

To achieve immunostaining, sections were washed three times in PBS (phosphate buffered saline, Hylabs) and then permeabilized in PBST (phosphate buffered saline containing 0.1% Triton X-100 (Merck)). Next, sections were blocked in PBST containing 20% NGS (normal goat serum (Vector Laboratories)) for 1 h at room temperature and incubated with primary antibodies in PBST (containing 2% NGS) at 4°C for 24-36 h. After 3 washes in PBS, sections were incubated with secondary antibodies conjugated to fluorophores in PBST containing 2% NGS for 1.5 h at room temperature. After three washes in PBST and once in PBS, the sections were mounted onto glass slides and cover-slipped with an aqueous mounting medium (Thermo Scientific, catalog # 9990412).

### Quantification and statistical analysis

#### *Sleep scoring*

We categorized different time periods to either active-wakefulness with behavioral activity (e.g., locomotion, grooming. Observable in video, in high EMG activity, and high frequency and low voltage EEG activity), quiet-wakefulness (low-voltage high-frequency EEG activity and high tonic EMG with occasional phasic activity), light NREM sleep (low to medium-amplitude slow wave EEG activity and low tonic EMG activity, a transitional state from wake to NREM), NREM sleep (high-amplitude slow wave EEG activity and low tonic EMG activity), NREM to REM transition (marked by especially high spindle density), REM sleep (low-amplitude wake-like frontal EEG co-occurring with theta activity in parietal EEG and flat EMG).
