## supplementary table 1 for "An early surge of norepinephrine along brainstem pathways drives sensory-evoked awakening"

| | Value | SE | tStat | pValue |
|---|---|---|---|---|
| **Intercept awakening** | **0.25** | **0.08** | **3.32** | **9.03 e-04** |
| **PRN surge awakening** | **-0.55** | **0.13** | **-4.11** | **4 e-05** |
| BF rise awakening | 0.02 | 0.19 | 0.1 | 0.92 |
