## Supplementary figure 1 for "An early surge of norepinephrine along brainstem pathways drives sensory-evoked awakening"

Figure S1: validating $GRAB_{NE}$ using optogenetics

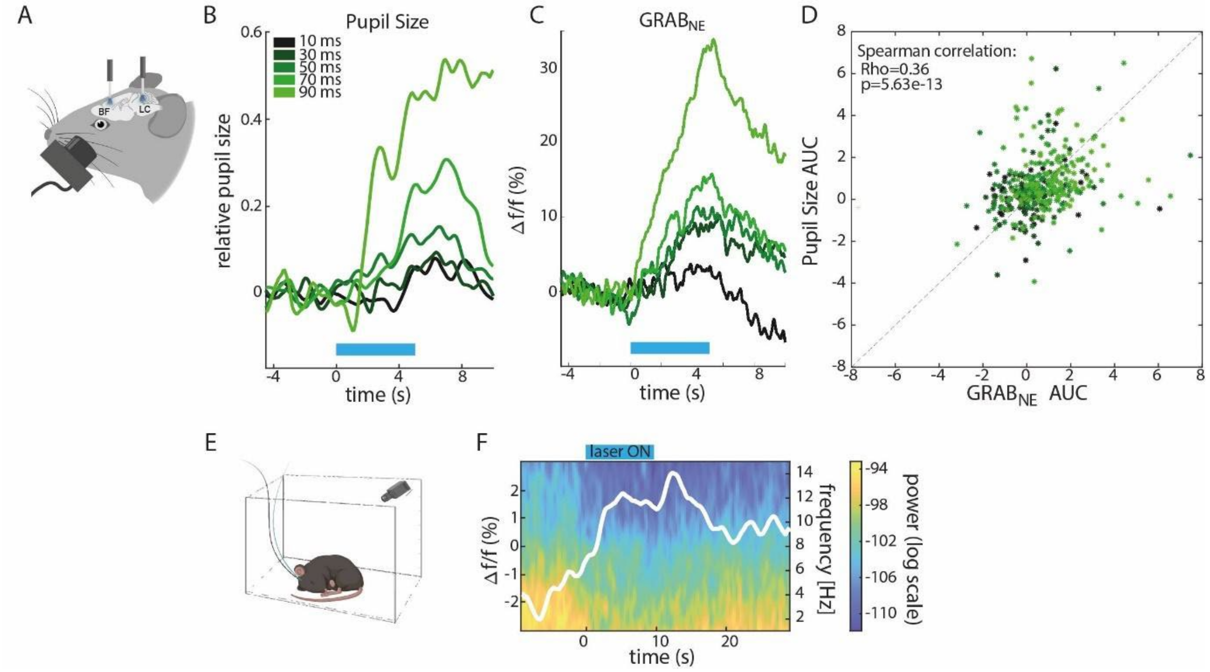


**A,** Depiction of surgical approach and experimental procedure. The photometry fiber was placed above the BF, while the LC was injected with CAV2-PRS-ChR2-mCherry and an optic fiber was placed above it (MFC_200/240-0.22_5mm_MF1.25_FLT, doric). Mice (n=5) were anesthetized using isoflurane and placed in the recording chamber on a heating pad. A camera (Logitech C615) was focused on the ipsilateral eye, an infrared light was placed to allow capturing the pupil in dark conditions and optic patch cords were connected. Laser stimulations (10mW) were automatically triggered every 30 seconds, randomly iterating between parameters. Duration was set to 5 secs, frequency to 10 Hz, and duty cycle [10, 30, 50, 70 or 90 ms]. Simultaneous video data were captured by a USB webcam with the IR filter removed, synchronized with photometry data. To extract pupil area we applied the same pipeline as in [14,38], video images were first cropped around the eye. A mask based on the median values was applied and the best centrally fitted circle was selected using the "regionprops" function in Mathworks MATLAB. For each trial separately, pupil area was normalized by the average baseline pupil area in the 5 second preceding trial onset, and percent change dynamics were calculated for the [-5 30] seconds interval around stimulation. Trials were then averaged for each animal separately (12 to 20 trials per mouse, n=5) to generate time-courses. **B,** Representative example of pupil dilation as consequence of laser activation. Lines represent the average pupil traces per stimulation intensity (from dark to light green) in a single mouse. The Cerulean horizontal line represents laser activation (t=0-5s). **C,** Representative example of $GRAB_{NE}$ signal in the BF as consequence of laser activation. Lines represent the average pupil traces per stimulation intensity (from dark to light green) in a single mouse. The Cerulean horizontal line represents laser activation (t=0-5s). **D,** Correlation of pupil size and $GRAB_{NE}$ area under the curve (AUC). Dots are trials across all mice. Spearman Rho=0.36, p=5.63e-13. **E,** Depiction of experimental setup of awakening experiment. **F**, Average spectrogram of EEG in a representative mouse across trials. White line represents average $GRAB_{NE}$ trace across trials. The Cerulean rectangle above represents laser activation (t=0-10 s).
