## Supplementary figure 2 for "An early surge of norepinephrine along brainstem pathways drives sensory-evoked awakening"

Figure S2: preprocessing and undisturbed sleep

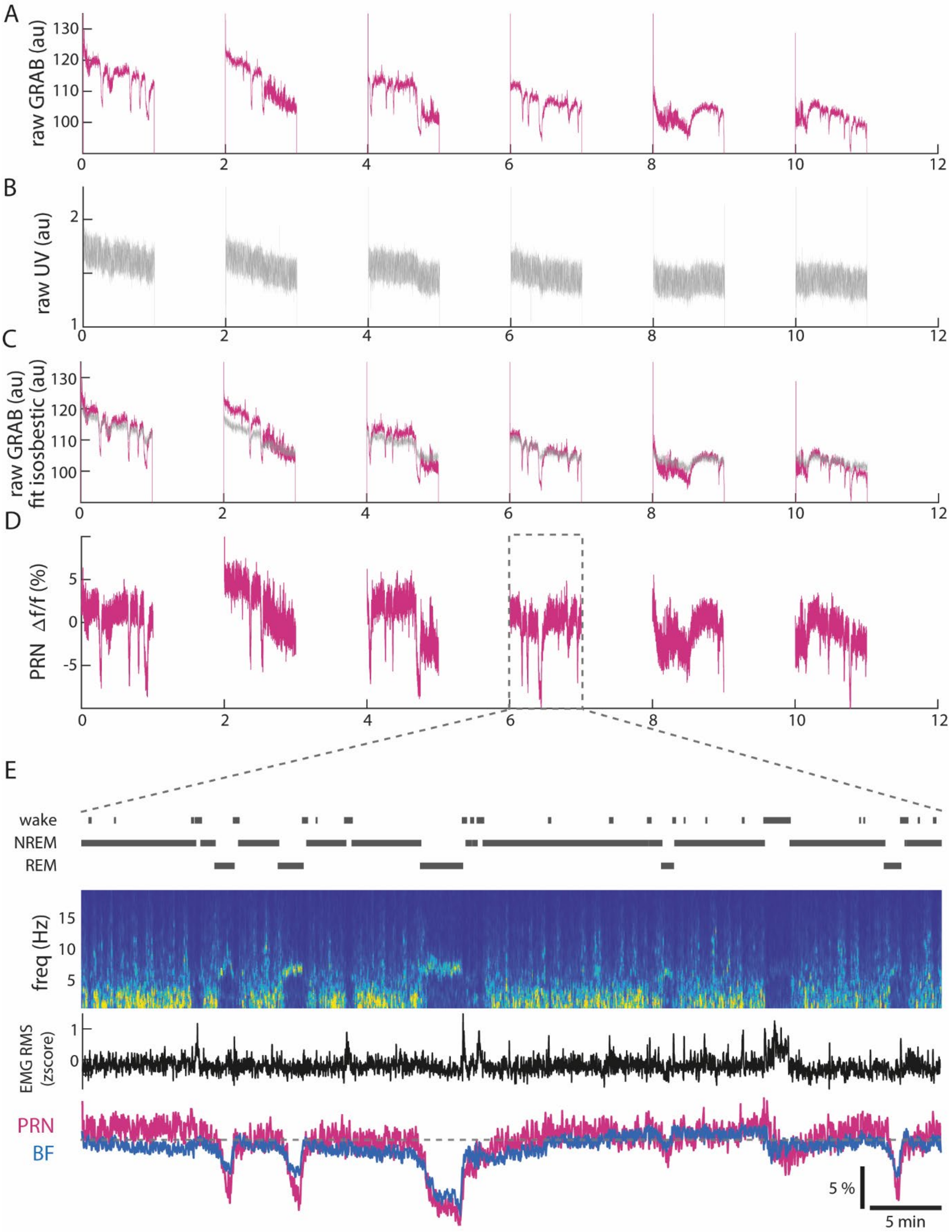


**A-D,** Representative example of preprocessing pipeline, as in[7]. **A,** Raw $GRAB_{NE}$ signal from PRN across a 12-hour recording. **B,** Raw isosbestic channel recording across 12 hours. **C**, Raw $GRAB_{NE}$ signal (magenta) overlaid with isosbestic channel after fitting (see methods). **D**, Δf/f= $\frac{GRAB - isosbestic}{isosbestoc}$ over 12 hours (see methods). **E**, One hour of full signal- from top to bottom: hypnogram, spectrogram of

EEG, EMG root mean square, $GRAB_{NE}$ signal from PRN and BF in magenta and blue accordingly. Dashed line represents 0%. Scale bars represent 5%- and 5-min.
