## Supplementary figure 3 for "An early surge of norepinephrine along brainstem pathways drives sensory-evoked awakening"

### Figure S3: retro GCaMP

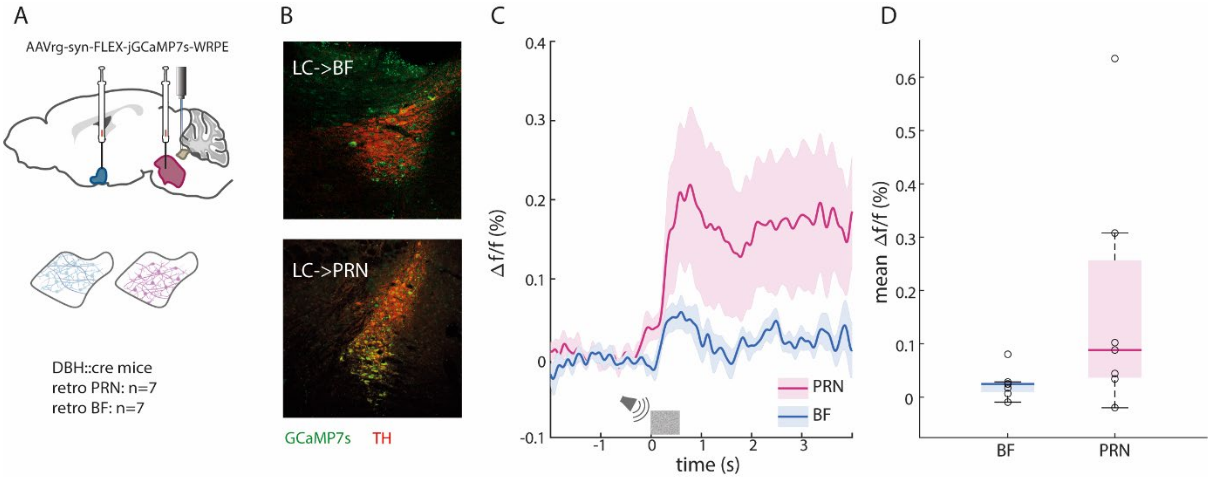


**A**, Depiction of surgical approach. A Cre-dependent retro AAV with GCaMP7s was injected to either PRN or BF (as described in Methods), and an optic fiber placed above the LC. **B**, Representative images of histology. **C,** Average GCaMP7s NREM SEA traces in LC->BF (blue) and LC->PRN (magenta) across mice ($n_{BF}$=7, $n_{PRN}$=7). **D,** Box plot of response averages (t=0.5-4s) in LC->BF and LC->PRN.
