## Supplementary figure 4 for "An early surge of norepinephrine along brainstem pathways drives sensory-evoked awakening"

Figure S4: Fluorophore only control for opto-activation

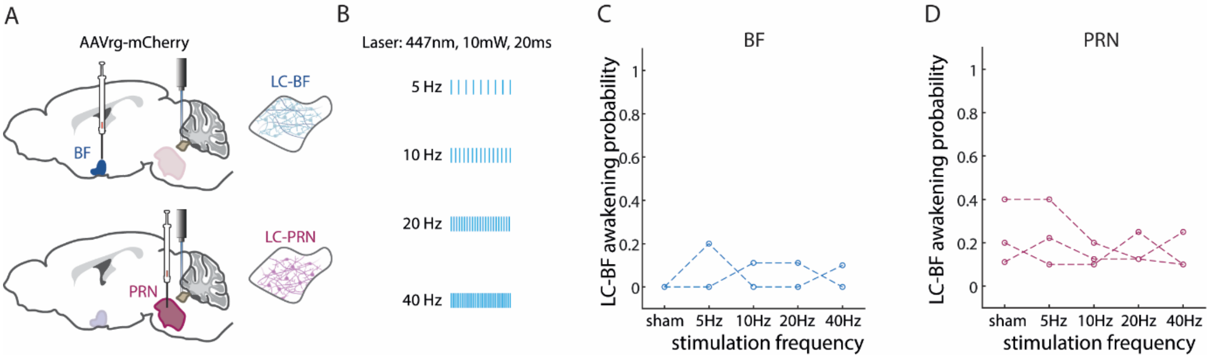


**A,** Depiction of surgical approach for LC->BF (top), and LC->PRN (bottom). **B,** Experimental procedure for laser awakening experiment. The experiment included 10s laser of 10mW and 20ms duty cycle for 5Hz, 10Hz, 20Hz and 40Hz. **C, D,** Probability to awaken from laser activation as a function of frequency for LC->BF mCherry (**C**) and LC->PRN mCherry (**D**). Dots represent single animals ($n_{BF}$=2, $n_{PRN}$=3)
