## Supplementary figure 5 for "An early surge of norepinephrine along brainstem pathways drives sensory-evoked awakening"

Figure S5: Fluorophore only control for selective synaptic silencing

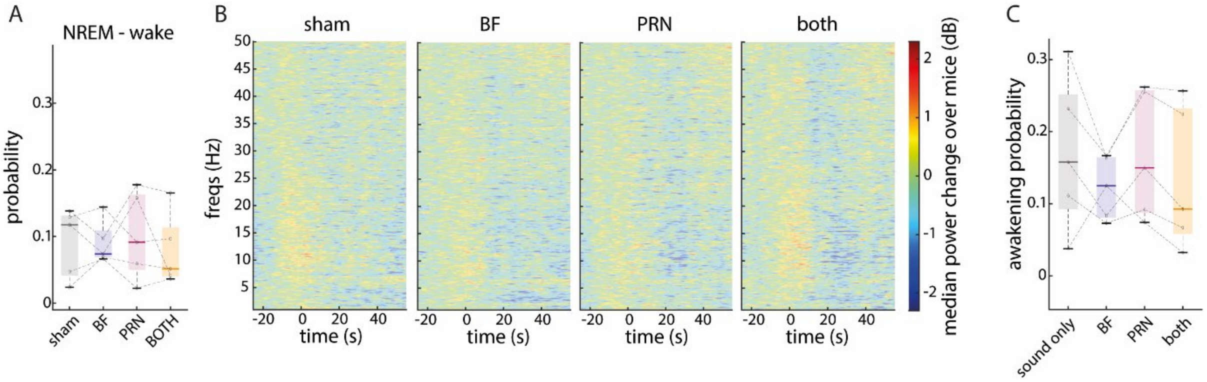


**A**, Box plot of average probability to transition from NREM to wake under each condition during t=5-10 from laser onset. Dots represents single animals (n=5). **B**, Median EEG spectrograms across mice around trials during NREM normalized to t=-30-0s relative to laser onset. Laser was on from t=0 to t=10s. left- sham condition, BF laser condition, PRN laser condition, right- both lasers on condition. Areas that are not significant compared to sham are opaque. **C**, Box plot of awakening probability from SEA. Gray- sham, blue- BF, magenta- PRN, yellow-both. Dots represents single animals (n=5).
